# Intranasal photobiomodulation as an energy-efficient, largely parameter-insensitive alternative to transcranial photobiomodulation

**DOI:** 10.64898/2026.08.08.743717

**Authors:** Alicia A. Mathew, Hannah Van Lankveld, Xiaole Z. Zhong, Joanna X. Chen, Reza Zomorrodi, J. Jean Chen

## Abstract

**Background:** Photobiomodulation (PBM) is an emerging non-invasive light-based brain stimulation technique that can alter cortical oscillations and is currently being pursued for improving cognition and treating neurological and psychiatric conditions. Nearly all human EEG evidence comes from transcranial PBM (tPBM) applied to the forehead, where light must traverse the scalp and thick skull, requiring protocols to compensate with high surface irradiance. Intranasal PBM (iPBM) can reach the anterior skull base at a fraction of that irradiance and has also been shown to modulate cerebrospinal fluid dynamics, yet it has been studied almost exclusively as an adjunct to tPBM, leaving its cortical effects in isolation, and its energy-efficiency relative to the transcranial route, unknown.

**Objective:** To define the spatiotemporal EEG response to pulsed iPBM delivered alone, determine whether stimulation parameters or individual biology moderate it, and compare the energy-efficiency of iPBM and tPBM in the same participants.

**Methods:** High-density EEG was collected from forty-six healthy young adults during pulsed iPBM and tPBM spanning a parameter space of varying wavelengths, pulsation frequencies, and irradiances. Percent change in band power from a within-session pre-stimulation baseline was tested with spatiotemporal cluster-based permutation tests. Linear mixed-effects models with backward elimination assessed stimulation and biological moderators (sex, nostril-to-cortex distance). Energy-efficiency, defined as the percent change in band power per J/cm^2^ of delivered surface energy, was compared between routes within each subject in delivery route-specific cluster regions of interest (ROI) (Wilcoxon signed-rank tests, Benjamini-Hochberg false discovery rate).

**Results:** iPBM alone produced significant spatiotemporal clusters in theta, beta, and gamma power, with anterior increases and posterior decreases; no delta or alpha clusters survived correction. Beta and gamma effects appeared at stimulation onset and persisted even after stimulation ended, whereas theta effects strengthened after stimulation ended. No predictor survived elimination in any band, time window, or cluster ROI: response magnitude was independent of wavelength, pulsation frequency, irradiance, sex, and nostril-to-cortex distance. Notably, although iPBM delivered roughly twenty times less surface energy than tPBM (∼0.6-1.1 vs ∼12-24 J/cm^2^), it produced EEG changes of similar magnitude, and its energy-efficiency exceeded that of tPBM in seven of eight eligible comparisons, with median iPBM-to-tPBM efficiency ratios of 14-32 (all FDR q<0.05)

**Conclusions:** Delivered in isolation, pulsed iPBM elicits a robust cortical EEG signature closely resembling that of tPBM, is insensitive to the stimulation parameters and individual factors tested, and achieves this at a small fraction of the delivered surface energy. As a result, delivery route, not surface irradiance alone, should be treated as a primary variable in PBM dose reporting and protocol design.

## 1 INTRODUCTION

Photobiomodulation (PBM) occupies a unique position among non-invasive brain stimulation techniques where, rather than inducing or applying currents to neural tissue, it delivers low-intensity near-infrared (NIR) light (∼600-1100 nm) that is absorbed by mitochondrial chromophores, primarily cytochrome c oxidase (CCO), promoting oxidative phosphorylation and ATP synthesis [1–3]. In humans, PBM has been associated with improvements in attention, memory [4,5], reaction time [6], and cognitive flexibility [7], and clinical trials are underway for neurodegenerative conditions, traumatic brain injury, stroke, and depression [8].

In order for PBM to advance into a precise, mechanistically grounded intervention, further investigation is required to establish how delivered light translates into measurable changes in brain activity. Electroencephalography (EEG) is well suited to this task as it indexes neural electrical activity at the scalp with high temporal resolution. Transcranial PBM (tPBM), in which light is applied to the scalp, has become the conventional route for delivering NIR light in humans, and most of what is known about its neural effects comes from EEG: PBM produces consistent shifts in cortical oscillations, most notably in the beta and gamma ranges implicated in cognition [9–11]. Transcranial delivery, however, requires light to traverse the scalp and thick skull. To compensate, tPBM protocols employ relatively high surface irradiance, yet both our own simulations and those of others indicate that only a small fraction of the applied energy reaches cortical tissue [12,13]. This attenuation motivates a different question of whether the brain can be reached along anatomical routes that place less tissue between the source and the target.

Intranasal PBM (iPBM) is one such route. Light delivered through the nasal cavity reaches the anterior skull base at the cribriform plate, adjacent to the orbitofrontal cortex and to nasal lymphatic and cerebrospinal fluid (CSF) pathways [14,15], and it does so at surface irradiances far below those used at the forehead. To date, however, iPBM has been studied almost exclusively as an adjunct to concurrent transcranial stimulation, and its cortical effects in isolation remain uncharacterized [9,14,15]. We recently showed that pulsed iPBM alone modulates CSF dynamics in healthy adults [16], using data from the same multimodal study reported here, in which the same cohort completed both EEG and magnetic resonance imaging (MRI) sessions. However, whether iPBM delivered alone produces measurable EEG responses, whether that response resembles the spatiotemporal profile established for tPBM, and whether it is shaped by stimulation parameters or individual anatomy still remain unknown.

Isolating iPBM also makes route comparison possible. Because the nasal and forehead delivery routes operate at surface irradiances that differ by more than an order of magnitude, the question instead becomes “how much neural change does each delivery route yield per unit of energy delivered?” Expressed in dose units, this defines an energy efficiency: percent change in band power per J/cm^2^ delivered, the quantity that dosimetry and protocol design ultimately turn on.

We, therefore, use high-density EEG to provide the first characterization of the cortical response to pulsed iPBM delivered without concurrent transcranial stimulation in healthy young adults. Our primary objective was to define the spatial and temporal signature of that response and to test whether it depends on stimulation parameters (wavelength, pulsation frequency, dose level) or biological factors (sex, nostril-to-cortex distance). Because every participant also received tPBM, our second objective was to compare the two routes on energy-efficiency within the same brains.

## 2 METHODS

### 2.1 Participants and nostril-cortex distance measurements

Forty-six healthy young adults (20-32 years; 24M/22F) were recruited from Baycrest’s participant database. Exclusion criteria included neurological or physiological disorders and substance use. Ethics approval was obtained from the Baycrest Research Ethics Board and all participants provided written informed consent. To account for inter-subject variability caused by anatomical differences in the nasal passage, the linear distance from the nostril to the cortex was measured with Freeview’s built-in measure tool (**Fig. 1**), using the subnasale (the intersection of the philtrum and the base of the nose), the angle of the passage, and the grey matter as landmarks in the right nostril passage.

**Figure 1.**
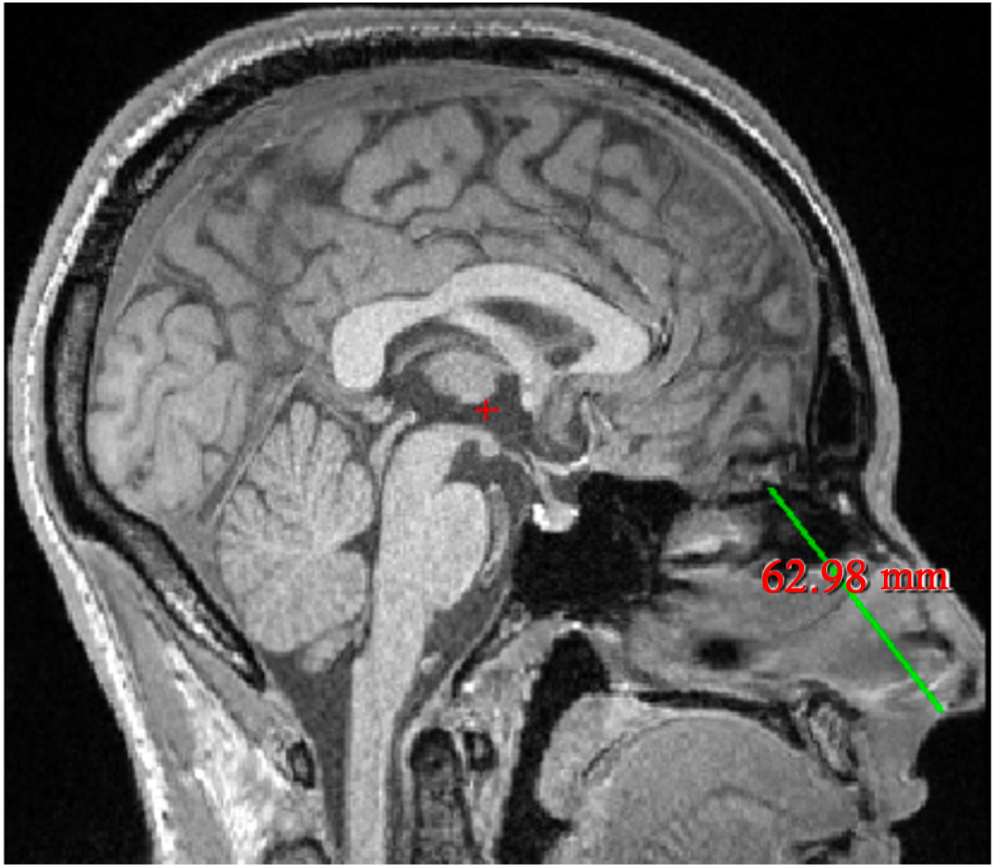
The distance between each participant’s nostril and cortex (in millimeters) with Freeview’s built-in measure tool.

### 2.2 Experimental design and iPBM protocol

Participants were randomly placed in one of three protocols which determined the laser parameters used in each of their four EEG-iPBM recordings (**Table 1**). These included two wavelengths (808 nm and 1064 nm), two pulsation frequencies (10 Hz and 40 Hz), and three irradiances (5, 7, and 9 mW/cm^2^). These were chosen based on their prevalence in PBM studies, their established tissue penetration and safety profiles, and their efficacy in past work [17]. The number of participants per protocol was as balanced as possible.

**Table 1.**
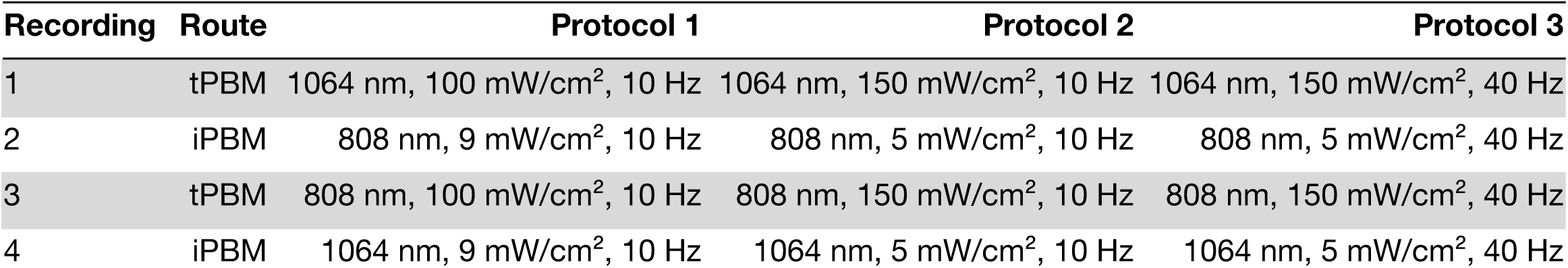
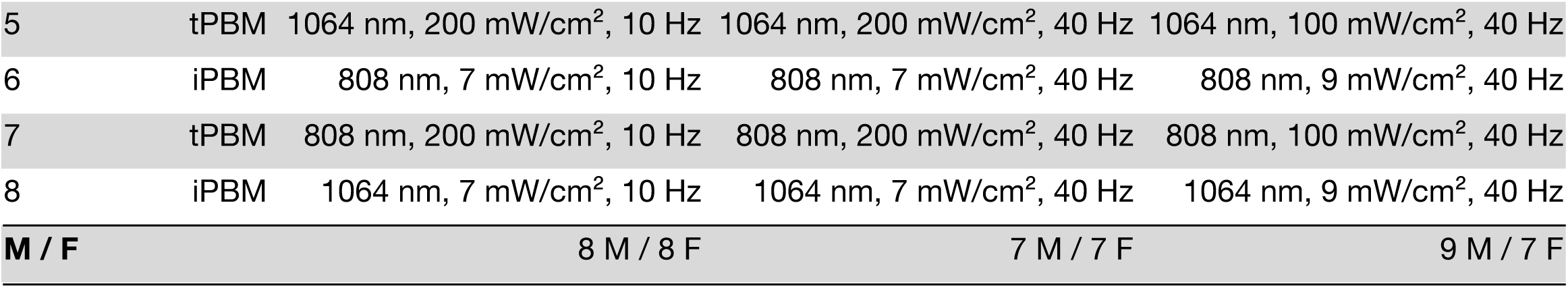
Participant distribution across protocols and stimulation parameters.

Each EEG recording was 12 minutes long and followed a PRE-DURING-POST block stimulus design (4 minutes each; **Fig. 2b**), during which participants sat and watched naturalistic stimulus videos to minimize drowsiness and control brain state [18] (**Fig. 2a**). In the absence of a formal sham condition, the PRE period served as a within-subject control or pre-stimulus baseline. In the DURING period, NIR light was administered through the nasal cavity via a nosepiece clipped to the right nostril, targeting the right prefrontal cortex (rPFC), using a 10-meter, 400 µm fiber cable and two Class 3 laser systems: MDL-III-808-1W and MDL-III-1064-1W (Vielight Inc., Toronto, Canada; **Figure 2a**) that delivered 808 nm or 1064 nm light, respectively. Laser parameters were regulated remotely so participants were blinded to the protocols and study design. Dosage parameters are in **Table 3**; calibration details in supplementary **Section S1**. Finally, to confirm the lack of measurable heating effects in the stimulated region caused by high irradiances during tPBM, a separate magnetic resonance (MR) thermometry scan was acquired, with participants also reporting no thermal sensations (supplementary **Section S2**).

**Figure 2.**
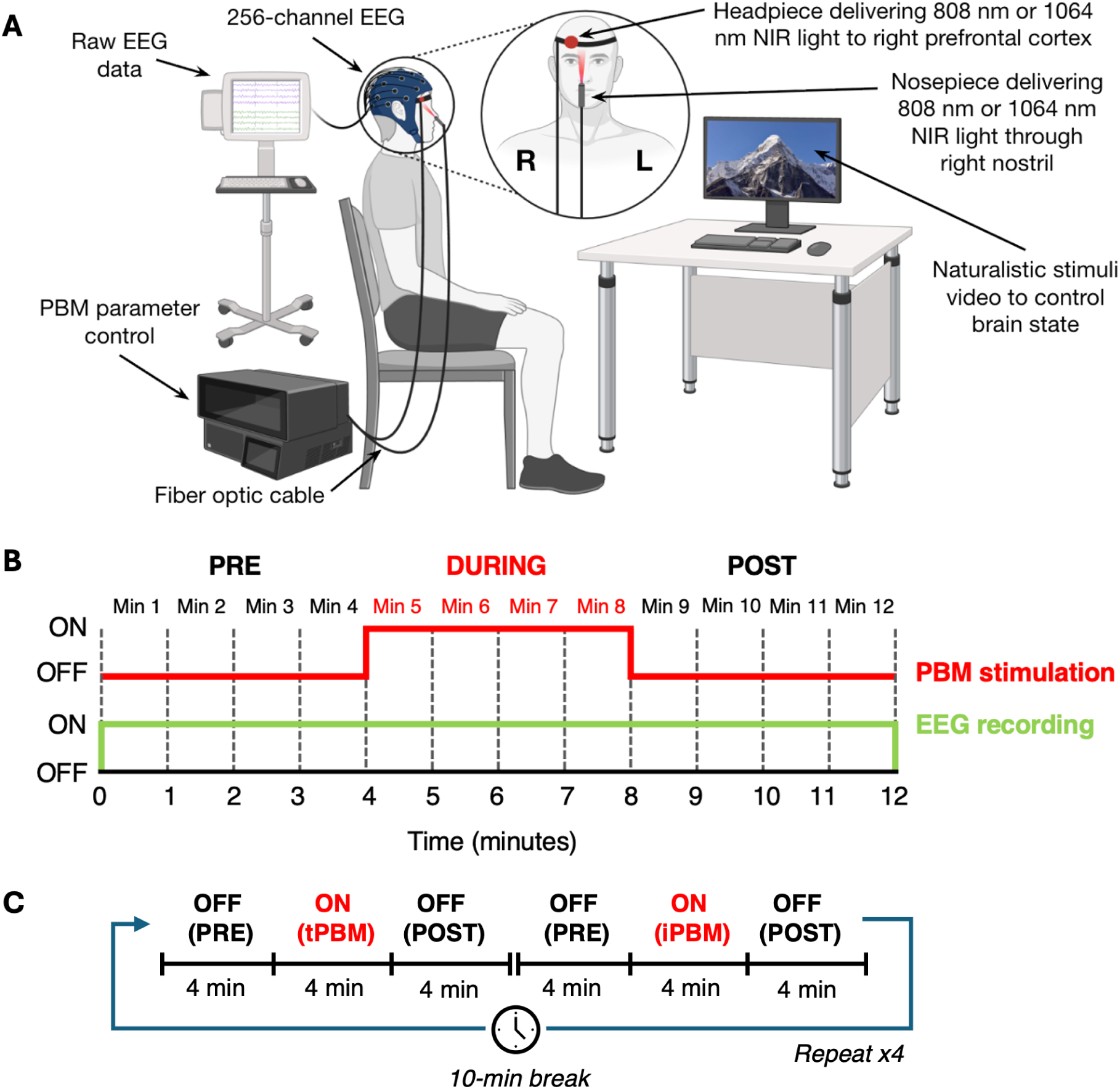
Experimental setup and recording timeline. (A) Schematic illustration of EEG-PBM experimental setup (*BioRender.com*). (B) EEG recording timeline following block PBM stimulus design. (C) Order of data acquisition: the iPBM block stimulus design took place immediately after tPBM, followed by a 10-minute break. This was repeated four times.

### 2.3 EEG acquisition

EEG was recorded at 1,000 Hz using a 256-channel HydroCel Geodesic Sensor Net with a NetAmps 400 amplifier (Magstim EGI, Eugene, OR, USA; saline-based; reference Cz; impedance < 50 kΩ; amplitude resolution 0.024 µV).

**Figure 3.**
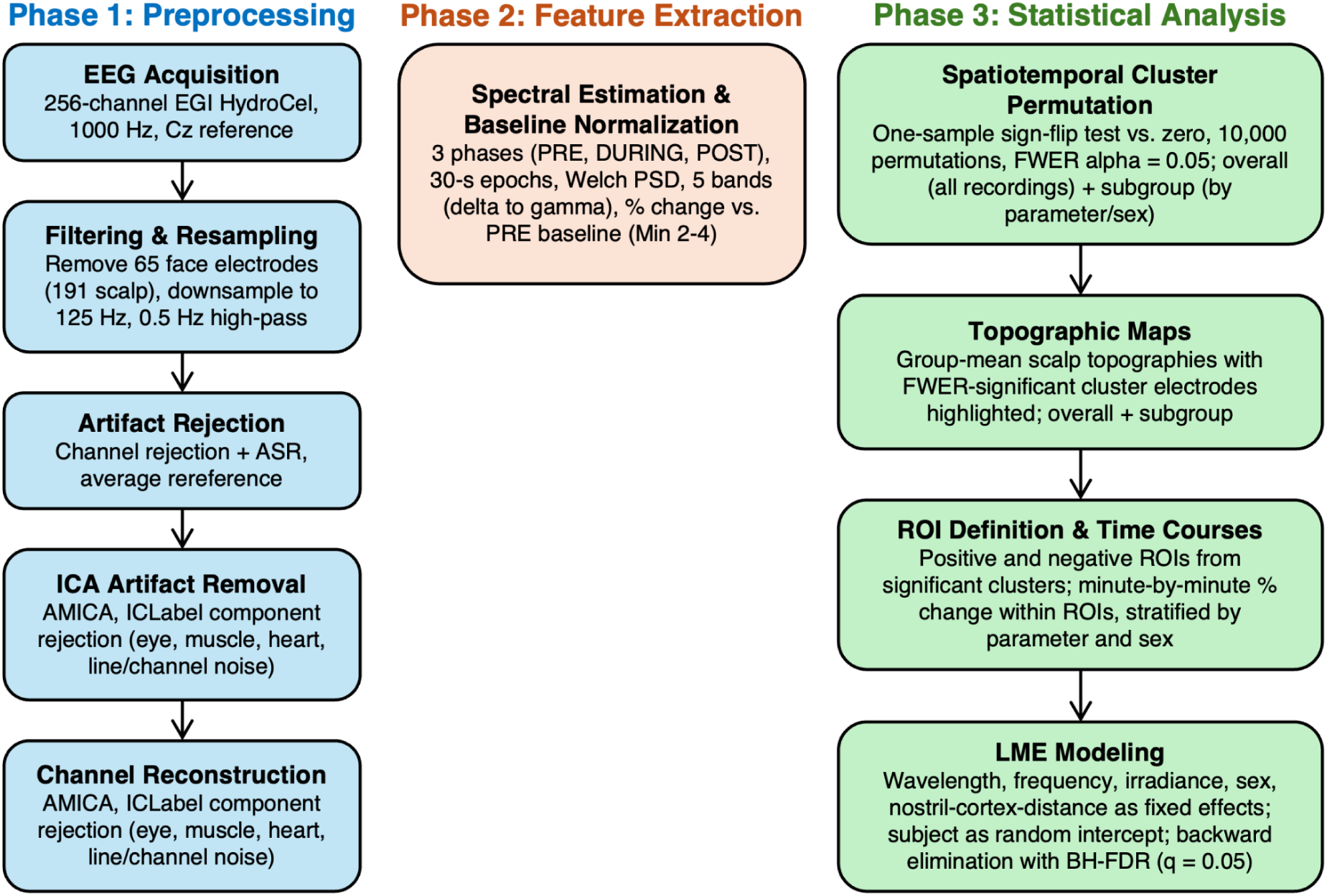
Summary of key stages in the data pipeline. EEG = electroencephalography; EGI = Electrical Geodesics Inc.; ASR = Artifact Subspace Reconstruction; ICA = Independent Component Analysis; AMICA = Adaptive Mixtures Independent Component Analysis; ICLabel = Independent Component Label; PSD = Power Spectral Density; FWER = Family-wise Error Rate; ROI = Region Of Interest; LME = Linear Mixed Effects; BH-FDR = Benjamini-Hochberg False Discovery Rate.

### 2.4 EEG data analysis

#### 2.4.1 Preprocessing

EEG data were preprocessed in MATLAB 2025b (MathWorks Inc., Natick, MA, USA) using EEGLAB 2026 [19]. 191 scalp channels were retained and data were downsampled to 125 Hz and high-pass filtered at 0.5 Hz (2nd-order zero-phase Butterworth). Noisy channels were then rejected automatically and remaining transients corrected using Artifact Subspace Reconstruction. Following average re-referencing, independent components were estimated with AMICA and those classified as muscle, eye, cardiac, line-noise, or channel-noise artifacts were removed. Rejected channels were then restored by spherical spline interpolation, a 55 Hz low-pass filter was applied after ICA, residual high-amplitude sample-level transients were interpolated, and recordings were trimmed to 720 s. Please see supplementary **Section S3** for additional details.

#### 2.4.2 Frequency band extraction and power calculation

Each electrode’s data was linearly detrended over the full recording and segmented into non-overlapping 30-second epochs. Power spectral density (PSD) was estimated via Welch’s method (2-second Hann windows, 50% overlap) with sub-window periodograms combined using a 20% trimmed mean. Band-limited absolute power was integrated within predefined ranges (delta: 1-4 Hz; theta: 4-8 Hz; alpha: 8-12 Hz; beta: 12-30 Hz; gamma: 30-50 Hz) and expressed as a percent change from the mean of the PRE baseline period (Min 2-4; **Eq. 2**). Minute 1 was excluded from the baseline due to instability.

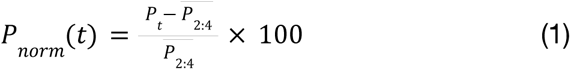

where P_t_ is the band power in minute t and 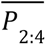 is the average band power over Min 2-4, and P_norm_ is the baseline-normalized absolute band power.

#### 2.4.3 Statistical analyses

##### 2.4.3.1 Spatiotemporal cluster-based permutation to identify significant electrode clusters

To identify changes in EEG frequency band power that were spatially and temporally consistent across subjects, cluster-based permutation tests [20] were performed using the MNE 1.12.1 package in Python 3.11.4. To study the overall effect of PBM as well as to observe any parameter-dependencies in the EEG responses, two group analyses were conducted. In the first analysis, all EEG recordings were averaged within-subject (**Fig. 4**) and in the second analysis, EEG recordings were first grouped by stimulation parameters or sex and then averaged within-subject (**Fig. 5**).

**Figure 4.**
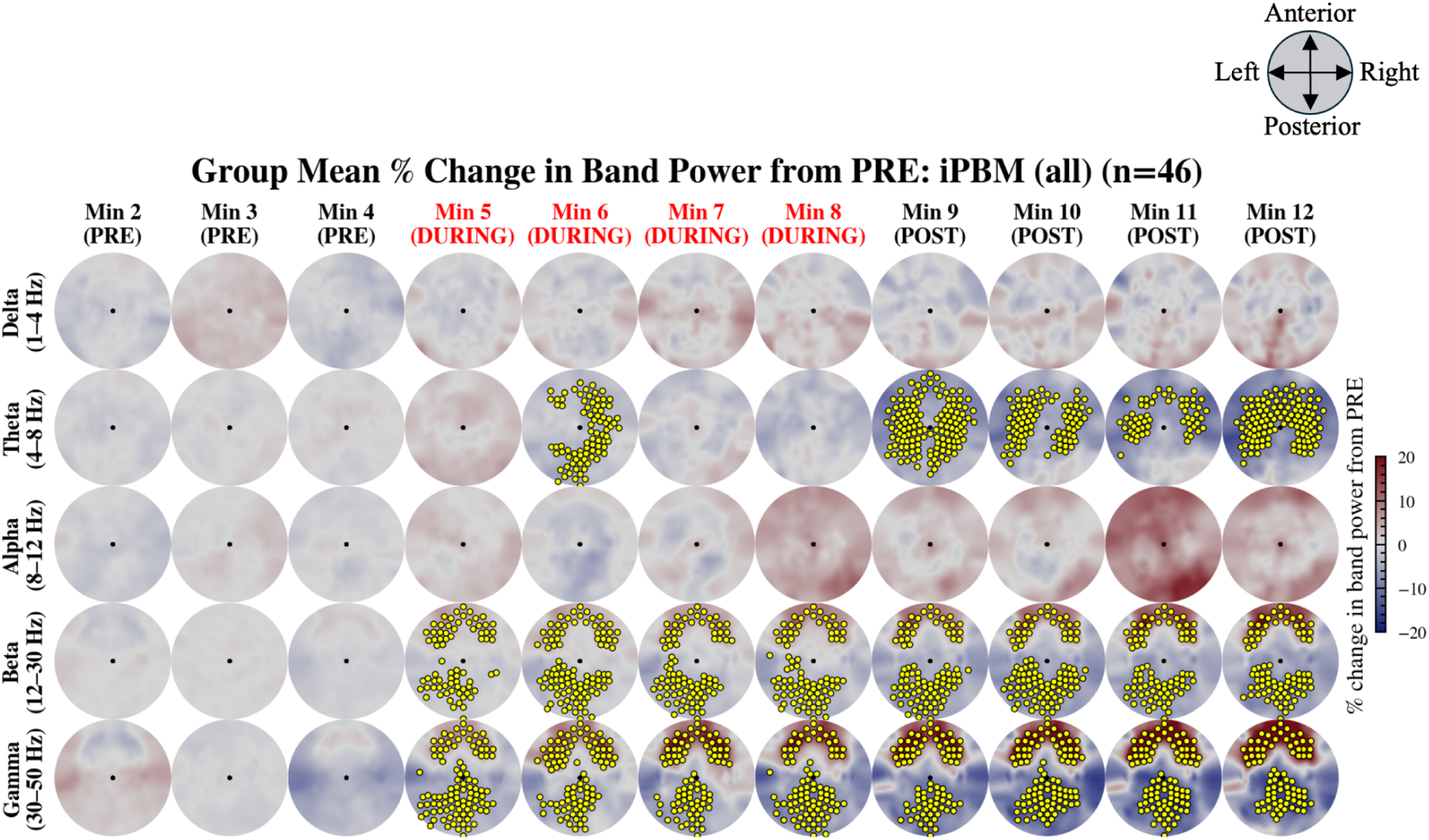
Group response to iPBM. Group-mean percent change in EEG band power from the mean of the baseline PRE period (Min 2-4) during iPBM (n=46). Rows: frequency bands (delta to gamma). Columns: successive 1-minute epochs (Min 2-12). Yellow markers: electrodes belonging to FWER-significant spatiotemporal clusters in that epoch (sign-flip permutation test, 10,000 permutations, cluster-level p<0.05). Epochs with iPBM stimulation (Min 5-8) are labeled in red. Color scale: ±20% change from PRE.

**Figure 5.**
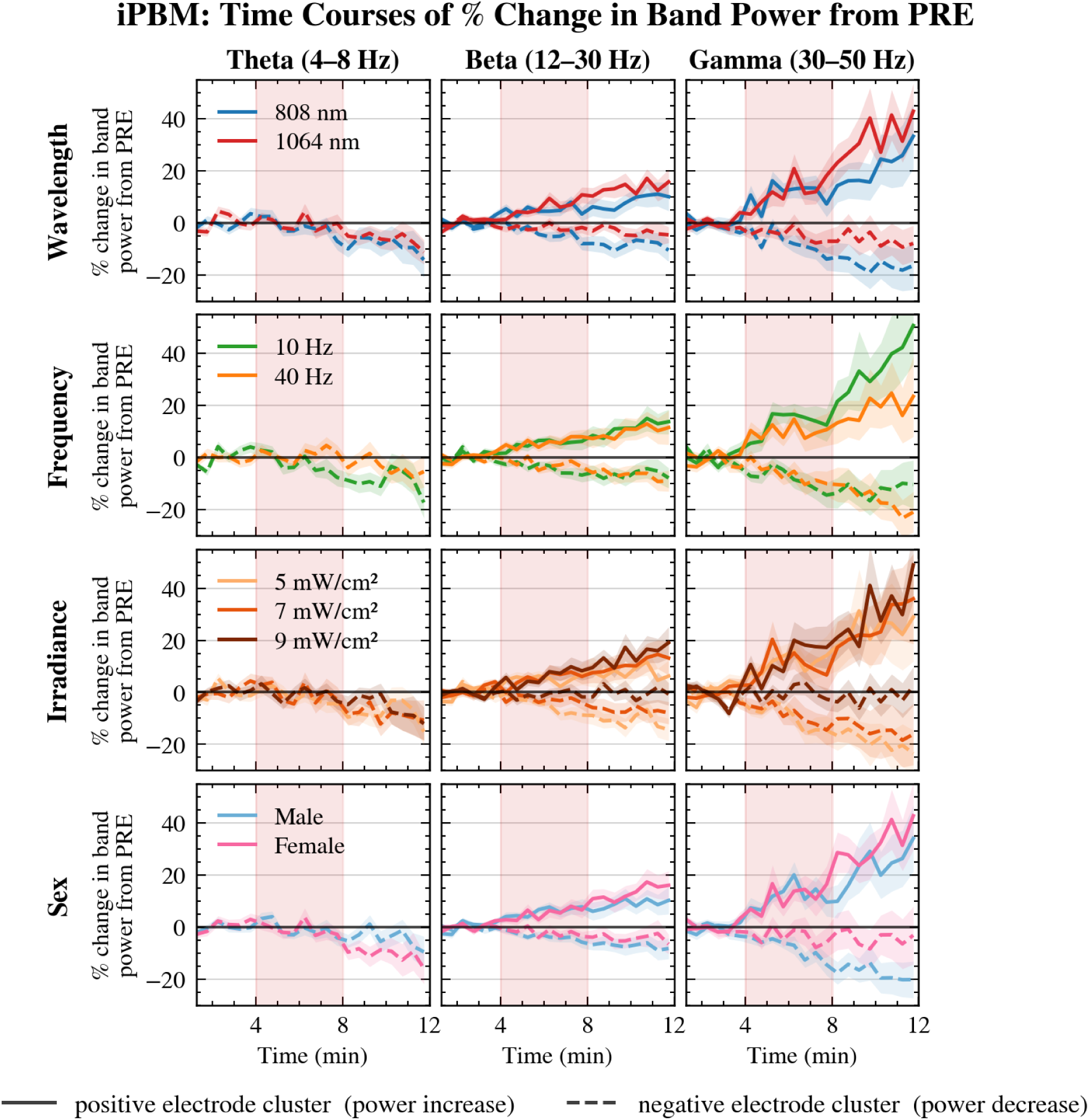
Group-level EEG band power responses to iPBM, stratified by stimulation parameters and sex. Minute-by-minute percent change in band power from the mean of the baseline PRE period (Min 2-4) is plotted within the FWER-significant positive (solid lines; power increase) and negative (dashed lines; power decrease) spatiotemporal cluster ROIs. Rows: parameter subgroups (wavelength, pulsation frequency, irradiance, sex); shaded bands indicate ±1 SEM across subjects. Columns: frequency bands with at least one significant cluster (theta to gamma). Red-shaded region: active iPBM stimulation period. Significant bar (solid) with asterisk: significant differences (FDR q<0.05) in parameter levels within the positive cluster ROI. Significant bar (dashed) with asterisk: significant differences (FDR q<0.05) in parameter levels within the negative cluster ROI. The position of the bars determine the time window over which the significant differences were present (during or post-stimulation) (see **Table 4**).

In each analysis, for each band x time window (DURING, POST), a 3D array (subjects ╳ time bins ╳ electrodes) was constructed where each time bin comprised 8 non-overlapping 30-second windows. Spatiotemporal adjacency was defined by combining temporal adjacency (consecutive windows) with electrode spatial adjacency (3D inter-electrode distances) via MNE’s combine_adjacency function. A one-sample sign-flip permutation test against zero was applied (two-tailed; 10,000 permutations). The cluster-forming threshold was an uncorrected alpha = 0.05 per time-electrode element. Cluster-level significance was controlled with a family-wise error rate (FWER) of alpha = 0.05 based on the maximum cluster-mass statistic across permutations.

##### 2.4.3.2 Time courses of significant electrodes

From the FWER-significant spatiotemporal clusters identified in **Section 2.4.3.1**, two regions of interest (ROIs) were defined: a positive ROI comprising electrodes belonging to any FWER-significant cluster with a positive group-mean change, and a negative ROI comprising those with a negative change. For each recording, percent-change values were averaged across ROI electrodes; recording-level time courses were then averaged within subjects; group means ±1 SEM were plotted per minute (**Fig. 8**). Only frequency bands with at least one FWER-significant cluster are shown. These same ROIs were used to visualize parameter-stratified time courses, permitting direct cross-group comparison on a common electrode set. No inferential tests were applied to time courses; parameter-level inference relied on mixed-effects models (**Section 2.4.3.3**).

##### 2.4.3.3 Modeling the effects of stimulation parameters and biological factors

Linear mixed-effects (LME) models were fit in Python (statsmodels v0.14; REML) to test the effects of stimulation parameters and individual biology on EEG responses within the FWER-significant electrode set. Models were fit separately for each frequency band, time window (DURING, POST), and responder class (positive ROI, negative ROI). The outcome Y was the recording-level mean percent change in band power averaged across the relevant ROI. Subject ID was included as a random intercept (experimental unit = recording session). The formula (Model A) is shown in **Eq. 2**:

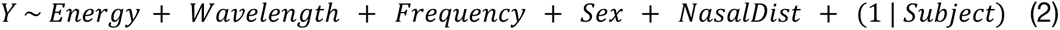

This formula is specific to the iPBM-EEG recordings as the tPBM-EEG analysis was done in our previous work. Categorical variables were Energy (Low: 5 mW/cm^2^, Mid: 7 mW/cm^2^, High: 9 mW/cm^2^), Wavelength (808 nm, 1064 nm), Frequency (10 Hz, 40 Hz), and Sex (Male, Female), while NasalDist was z-scored and included as a continuous variable. The reference levels were Low (Energy), 808 nm (Wavelength), 10 Hz (Frequency), and Male (Sex). Backward elimination was applied: at each step, Benjamini-Hochberg false discovery rate (FDR) correction (q=0.05) was applied simultaneously to all fixed-effect p-values, and the predictor with the highest FDR-corrected q-value was removed if it exceeded 0.05. Elimination continued until all contrasts of all remaining predictors were FDR-significant.

Model A coefficients are provided in supplementary **Table S2**, and the backward elimination trace is provided in supplementary **Table S3**. A separate carry-over analysis (Model B) tested whether the energy level of the immediately preceding recording session predicted the current response (see supplementary **Table S4**).

##### 2.4.3.4 Quantifying energy-efficiency

To investigate whether iPBM produces more EEG band power change per unit of delivered energy than tPBM, both modalities (iPBM, tPBM) were compared within each subject and the contrast was summarized at the group level. Efficiency was computed within each modality’s own FWER-significant spatiotemporal cluster ROI (**Section 2.4.3**).

Total delivered energy (fluence; J/cm^2^; **Table 2**) was calculated using **Eq. 3**. For each recording, band, and phase (DURING, POST), efficiency (**Eq. 4**) was the mean percent change across ROI electrodes divided by the delivered energy during that EEG recording:

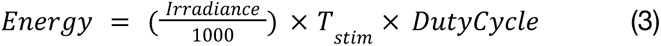

**Table 2.** PBM dosage parameters.

| Parameter | tPBM Value | iPBM Value |
| --- | --- | --- |
| Beam spot size at stimulation site (cm<sup>2</sup>) | ~1 | ~1 |
| Average distance from scalp (cm) | ~0 | - |
| Average distance from nasal laser to brain (mm) | - | 65.3 |
| Wavelength (nm) | 808, 1064 | 808, 1064 |
| Pulsation frequency (Hz) | 10, 40 | 10, 40 |
| Duty cycle (%) | 50 | 50 |
| Peak irradiance at stimulation site (mW/cm<sup>2</sup>) | 100, 150, 200 | 5, 7, 9 |
| Average irradiance at stimulation site (mW/cm<sup>2</sup>) | 50, 75, 100 | 2.5, 3.5, 4.5 |
| Duration of stimulation (minutes) | 4 | 4 |
| Total energy exposure at stimulation site (J) | 12, 18, 24 | 0.6, 0.84, 1.08 |
| Total energy density at stimulation site (fluence) (J/cm<sup>2</sup>) | 12, 18, 24 | 0.6, 0.84, 1.08 |

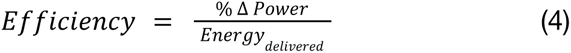

Cluster electrodes were split into positive and negative sub-ROIs by the sign of the group-mean percent change. A band ╳ phase ╳ ROI-direction comparison was included only when both modalities had at least three electrodes in that sub-ROI. For each such contrast, each subject’s efficiency was averaged across their four recordings per modality, and tPBM versus iPBM efficiency was tested with a paired Wilcoxon signed-rank test (n=46). P-values were FDR-corrected across the set of eligible contrasts (supplementary **Table S5**; **Fig. 6**).

**Figure 6.**
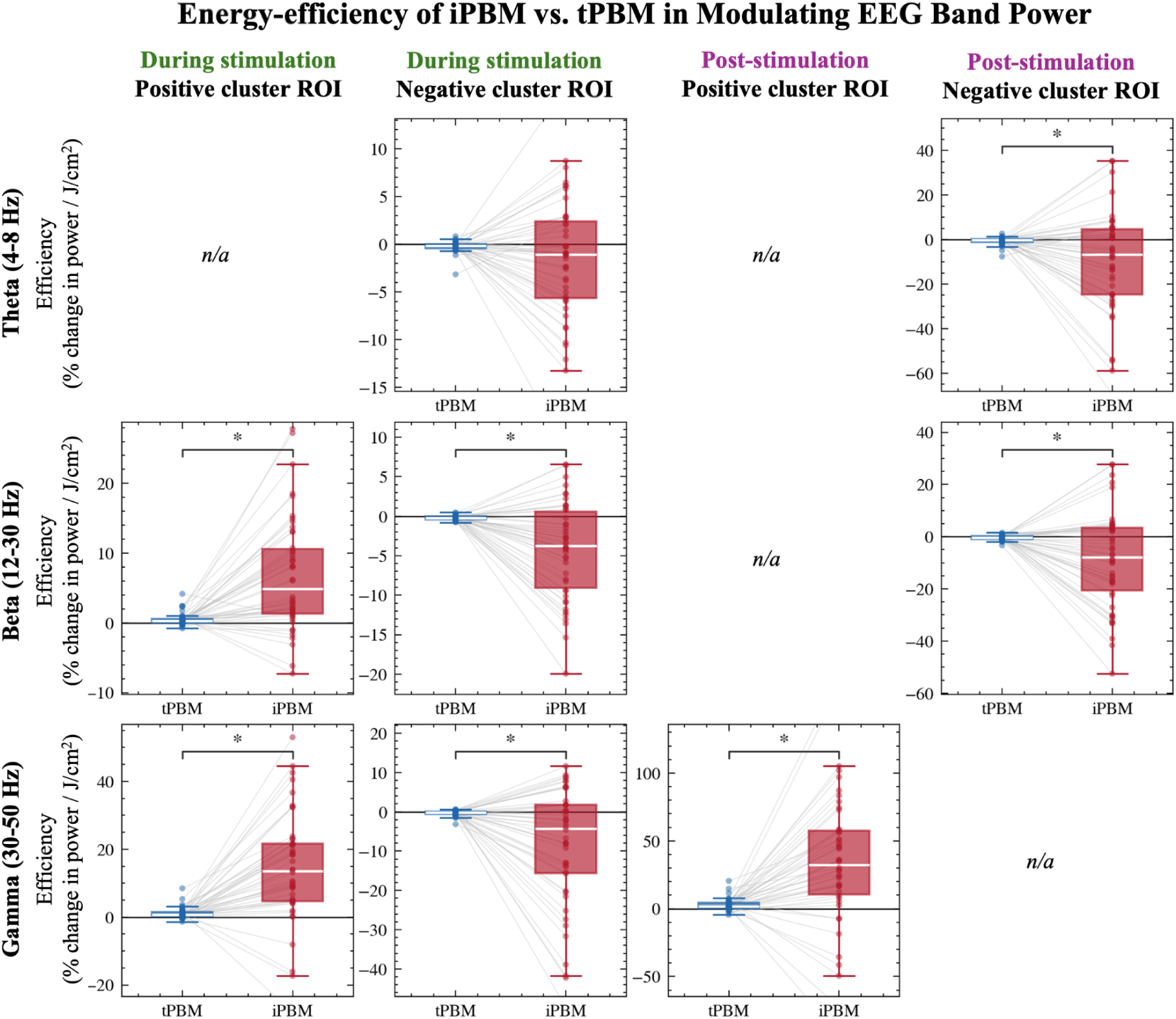
Energy efficiency of iPBM versus tPBM. Paired subject-level efficiency (% change in band power per J/cm^2^ delivered) for tPBM (blue) and iPBM (red) within each modality’s FWER-significant cluster ROI. Rows are frequency bands with eligible joint comparisons (theta, beta, gamma); columns are time window × ROI direction (During/Post × positive-/negative-cluster electrodes). Each panel shows subject averages across recordings (n=46), with boxplots, individual points, and within-subject links. Asterisks mark FDR-significant paired Wilcoxon contrasts (q<0.05; supplementary **Table S5**). Panels labeled “n/a” were not tested because both modalities did not each have ≥3 electrodes in that sub-ROI.

## 3 RESULTS

### 3.1 Spatiotemporal EEG response to pulsed iPBM

**Fig. 4** shows group-mean scalp topographies for all 46 participants across recorded minutes. No significant spatiotemporal clusters were identified for delta or alpha power. Theta power showed significant anterior and posterior modulations that were small in the second minute of stimulation (Min 6) and became more widespread and negative through the post-stimulation period (Min 9-12). Beta and gamma power were significantly modulated from stimulation onset (Min 5) through the end of the recording (Min 12). Significant beta and gamma increases were concentrated frontally/anteriorly, whereas significant decreases were observed posteriorly, with gamma responses larger in magnitude than beta.

### 3.2 Parameter and biological dependencies in the spatiotemporal EEG response to pulsed iPBM

**Fig. 5** shows minute-by-minute percent change from the pre-stimulus baseline, stratified by stimulation parameters and sex. Consistent with the absence of clusters in **Section 3.1**, delta and alpha power were not significantly modulated by any parameter or by sex. For bands with significant iPBM effects, LME analyses revealed no significant effects of wavelength, pulsation frequency, irradiance, sex, or nostril-to-cortex distance on response magnitude (supplementary **Table S2**). Across stratified time courses, responses tended to grow in magnitude after stimulation ended.

### 3.3 Energy-efficiency of iPBM versus tPBM

**Fig. 6** compares subject-level energy efficiency (percent change in band power per J/cm^2^ delivered) between iPBM and tPBM within modality-specific FWER cluster ROIs. In seven of eight eligible contrasts, iPBM was significantly more efficient than tPBM (paired Wilcoxon signed-rank tests, BH-FDR q<0.05; supplementary **Table S5**). Where power increased, iPBM was about 14-16 times more efficient than tPBM (beta, during stimulation; gamma, during and after stimulation). Where power decreased, iPBM was about 17-32 times more efficient (beta, during and after stimulation; gamma, during stimulation; theta, after stimulation). Overall,iPBM produced substantially larger EEG band-power change per unit delivered energy than tPBM in the shared, statistically eligible cluster ROIs.

## 4 DISCUSSION

In this work, we found that iPBM (i) significantly modulates power in largely the same frequency bands and directions as tPBM (theta-gamma), with the exception of alpha; (ii) was not significantly influenced by the stimulation parameters or biological factors tested here; and (iii) was more than an order of magnitude more efficient than tPBM at eliciting these power changes, despite delivering roughly one-twentieth of the surface energy per recording. Below we interpret these findings and discuss their implications for dose reporting and protocol design.

### 4.1 Pulsed intranasal PBM engages the cortex similar to pulsed transcranial PBM

The spatiotemporal profile of the iPBM-EEG response – progressive theta suppression and frontally dominant beta/gamma increases that strengthened after stimulation – parallels the shift from lower- to higher-frequency activity that has become a hallmark of tPBM in healthy adults [9–11]. The emergence of this pattern when light was delivered through the nasal cavity alone indicates that intranasal delivery is sufficient to drive cortical oscillatory change, rather than acting only as an adjunct to forehead stimulation. Frontal predominance of high-frequency effects is consistent with the anatomical trajectory of intranasal light near the cribriform plate and orbitofrontal cortex. The delayed build-up into the post-stimulation window further mirrors tPBM and is compatible with metabolic and vascular cascades that continue after illumination ends [21,22]. The absence of significant alpha clusters for iPBM, despite alpha effects under tPBM in related work, is also noteworthy. This may reflect true route-dependent differences in network engagement, reduced sensitivity for alpha under the present iPBM dose range, or simply lower spatial/temporal consistency and larger inter-subject variability of alpha effects after multiple-comparison control. Overall, these EEG findings complement prior work in the same cohort showing iPBM’s ability to also modulate cerebrospinal fluid dynamics [16].

### 4.2 EEG response to pulsed iPBM was largely parameter-insensitive

LME analyses revealed no significant effects of iPBM wavelength, pulsation frequency, irradiance, sex, or nostril-to-cortex distance on EEG response magnitude. This contrasts with tPBM in the same cohort, where wavelength and pulsation frequency were strong moderators of response magnitude, with 808 versus 1064 nm associated with differing spatial profiles and 10 versus 40 Hz differentially engaging gamma power, and where sex was a prominent biological moderator. The absence of analogous parameter and sex effects for iPBM suggests that, within the tested range, intranasal cortical EEG responses may be more predictable across dose parameters than forehead responses, while still remaining large enough for a clear efficiency advantage over tPBM.

### 4.3 Intranasal is markedly more energy-efficient than transcranial delivery

When EEG responses were normalized by delivered surface energy, iPBM outperformed tPBM by roughly an order of magnitude (median efficiency ratios ∼14-32 times) across eligible theta, beta, and gamma contrasts. Group topographies show that absolute percent-change magnitudes for the shared responses, particularly the frontally positive and posteriorly negative beta and gamma patterns, are in the same range for both routes, with tPBM appearing only modestly larger, despite delivering ∼20 times more surface energy (∼12-24 versus ∼0.6-1.1 J/cm^2^). Under that condition, energy normalization is the appropriate way to express that comparable scalp EEG neuromodulation was obtained at a fraction of the optical cost. Efficiency here refers to EEG change per unit delivered surface energy, not per photon absorbed at cortex thus, whether the gap reflects higher cortical absorption along the shorter intranasal path, differences in engaged pathways, or both will require route-specific dosimetry. Practically, however, the implication is clear: for these oscillatory endpoints, intranasal delivery achieves similar effects at much lower delivered energy than forehead stimulation.

### 4.4 Delivery route as a core variable in precision neuromodulation

These findings argue that, in PBM, delivery route belongs in the dose equation alongside wavelength, pulse structure, and irradiance. Surface irradiance (mW/cm^2^) is routinely used to label protocols, yet it is poorly suited for cross-route comparison: intranasal and forehead protocols with very different irradiance labels can produce related neural signatures, and only energy normalization revealed the large efficiency difference between them. For PBM to advance as a precision neuromodulation tool, studies should report delivered energy (J/cm^2^), specify route explicitly, and, where possible, report neural effect per joule. Irradiance-only reporting cannot support meaningful cross-study comparison or rational optimization across delivery routes.

### 4.5 Limitations and future work

Several limitations shared with our tPBM work in this cohort also apply here. The sample comprised healthy young adults, so EEG responses may differ in clinical populations with mitochondrial or network dysfunction. The repeated-recording design enabled within-participant comparisons but leaves residual risk of order effects: although carry-over screening uncovered an effect of the prior tPBM energy in only one iPBM outcome (gamma decreases during stimulation; supplementary **Table S4**), fatigue, habituation, and expectation cannot be fully excluded without greater temporal separation, more systematic parameter counterbalancing, or dedicated sham arms for every parameter combination. As before, the within-session PRE baseline is sufficient to detect stimulation-locked change but cannot explicitly rule out expectancy. The 12-minute window may also truncate the post-stimulation effects, given that several bands were still evolving at the final time point. We did not collect concurrent behavioral or cognitive measures, so the oscillatory changes cannot be linked directly to function.

Limitations more specific to the present iPBM analyses also warrant note. Efficiency was defined as EEG percent change per unit delivered surface energy within modality-specific cluster ROIs; this quantifies practical neuromodulation cost for scalp endpoints, but does not estimate absorbed cortical dose, and route-specific optical modeling will be required before the efficiency gap can be attributed solely to higher target energy deposition along the shorter intranasal path. The tested iPBM parameter space also remains finite, and the incomplete irradiance design limits formal within-route dose-response inference for energy-normalized outcomes. Finally, because these analyses remain at the scalp, they cannot localize the true sources contributing to cortical electrical activity or establish whether iPBM preferentially engages deeper structures along its optical pathway, paving the way for future focused work.

Complementary fMRI evidence in this cohort already shows that iPBM can modulate subcortical structures [23] therefore, EEG source reconstruction in the same paradigm would help reconcile cortical oscillatory effects with those deeper targets. A key next step is to integrate EEG with autonomic, and cerebrospinal fluid measures to link band-power change to other physiological pathways, including autonomic tone. Such work would clarify which components of the iPBM response are cortical, which are subcortical or systemic, and how delivery route shapes this broader physiological cascade.

## 5 CONCLUSION

Pulsed intranasal PBM produces a robust scalp EEG response that closely resembles the spectral signatures elicited by forehead tPBM, most notably theta power suppression and frontal beta/gamma increases that strengthen after stimulation, while remaining largely insensitive, within the tested range, to stimulation parameters and individual biology. When responses were normalized by delivered surface energy, iPBM emerged as more than an order of magnitude more efficient than tPBM, achieving substantial neuromodulation at a small fraction of the fluence. These results position intranasal delivery as a promising low-exposure, safe, and potentially more convenient route for PBM treatment and establish delivery route as a variable that must be measured, reported, and optimized alongside conventional optical dose parameters.

## 6 DATA AVAILABILITY

Data and software code can be made available upon request.

