## Supplementary Materials for "Intranasal photobiomodulation as an energy-efficient, largely parameter-insensitive alternative to transcranial photobiomodulation"

### S1 Stimulation calibration and dosimetry

#### S1.1 Optical power calibration

Prior to data acquisition, laser output was measured and calibrated using a Newport optical power meter (Model 843-R, Newport Corporation, USA). The optical fibre was connected to the laser power supply on one end and the forehead applicator on the other. The applicator was placed directly on the power meter sensor to ensure that the measured power reflected the delivered output to the skin surface.

The signal generator was connected to the power supply using a BNC cable. The laser mode was set to modulation, and the signal generator was configured to the target pulsation frequency (10 Hz or 40 Hz) with a 50% duty cycle and 5V amplitude.

#### S1.2 Energy and irradiance dose calculation

The applied irradiance was calculated by dividing the measured optical power (mW) by the cross-sectional area of the beam at the applicator tip, reflecting the time-averaged power across pulse cycles. The beam diameter was approximately 11.28 mm, corresponding to a radius of 0.565 cm and an overall area of 1cm$^2$ ($A = \pi * r^2$). Delivered energy (J/cm$^2$) was calculated as the product of applied irradiance (mW/cm$^2$), duty cycle and stimulation duration (240 s).

The applied irradiance was calculated by dividing the measured time-averaged optical power (mW) by the cross-sectional area of the beam at the applicator tip. The beam diameter was approximately 11.28 mm, corresponding to a radius of 0.565 cm and an area of approximately 1 cm$^2$ ($A = \pi * r^2$). Delivered energy density (J/cm$^2$) was calculated as the product of applied irradiance (mW/cm$^2$), duty cycle, and stimulation duration (240 s).

### S2 MR Thermometry

To evaluate potential temperature changes near the site of irradiation during tPBM, MR thermometry data were collected at the highest irradiance condition (1064 nm, 200 mW/cm$^2$) for both pulsation frequencies (10 Hz and 40 Hz) across 30 subjects. As the tPBM laser was positioned over the right forehead to stimulate the right prefrontal cortex (rPFC), this region served as the target region of interest (ROI) for thermometry measurements (**Fig. S1**). Group-level statistical testing was performed using paired-sample t-tests for each pulsation frequency, comparing the mean temperature during the baseline period (Min 1-4) to the stimulation period (Min 4-8).

As shown in **Table S1**, the average temperature change was minimal and remained stable over the entire 12-minute recording. This confirmed that even at the longest wavelength and highest irradiance, no measurable heating effects were produced in the targeted brain region. As the lasers were secured such that participants were unable to see any light, and no thermal sensations were reported, both thermal and placebo effects were minimized.

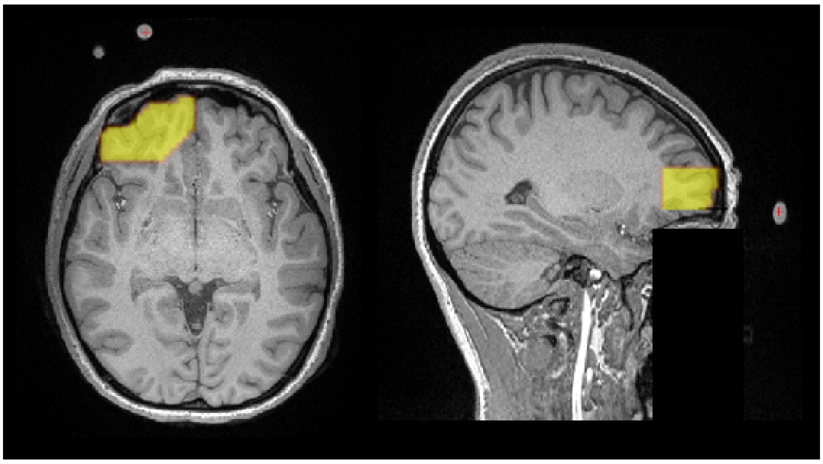

**Figure S1. The region of interest for thermometry assessment.** Vitamin E capsules were used as fiducial markers to indicate the laser location. The illuminated region encompasses approximately 3.59 cm from the site of light irradiation. This region was segmented for each participant and used for MR thermometry analysis.

**Table S1.** Group-level statistical comparison of temperature changes between the baseline and tPBM stimulation periods for both pulsation frequencies.

| Frequency | Baseline (°C) | tPBM (°C) | p |
|---|---|---|---|
| 10 Hz | -0.016 | 0.050 | 0.13 |
| 40 Hz | -0.0020 | 0.0087 | 0.76 |

### S3 EEG data preprocessing

A zero-phase second-order Butterworth high-pass filter (0.5 Hz cutoff) was applied to remove slow drifts while preserving delta-band oscillations. Automated channel rejection and Artifact Subspace Reconstruction (ASR; channel correlation = 0.85; flat-line = 5 s; line-noise = 4; burst criterion = 10 SD) were applied via EEGLAB's clean_rawdata; epoch rejection was disabled. An average reference was computed before Independent Component Analysis (ICA), which was performed using Adaptive Mixture ICA (AMICA)45 (one mixture; max 2000 iterations; PCA dimensionality = n-1). ICLabel was used to classify components; those labeled as Muscle, Heart, Line Noise, or Channel Noise (≥0.80) or Eye (≥0.70) were removed. Rejected channels were restored by spherical spline interpolation, and residual spikes exceeding twenty times the channel's robust noise level above 300 μV were linearly interpolated.

### S4 Model A: LME model coefficients

**Table S2. Summary of the primary LME model (Model A) for iPBM-induced EEG band power changes.** Each row represents one frequency band, time window (During-iPBM or Post-iPBM), and cluster ROI (Increase or Decrease) combination for which FWER-significant spatiotemporal clusters were identified in **Section 2.4.3.1**. All models began with the same five fixed-effect predictors (Energy, Wavelength, Frequency, Sex, and NasalDistance) and a random intercept for the subject. Backward elimination with Benjamini-Hochberg FDR correction (q=0.05) was applied iteratively (see **Table S3** for the full elimination trace). "Surviving Predictors" lists the predictors retained in the final model; a blank indicates that all predictors were eliminated. N Electrodes denotes the number of electrodes in the cluster ROI used as the dependent variable. "Estimate" reflects the mean difference in percent change from the mean of the PRE period relative to the reference level (808 nm, 10 Hz, Low energy, Male, high NasalDistance). Asterisks denote FDR-significant contrasts (q<0.05).

| Time Window | Band | ROI | N Electrodes | Initial Predictors | Surviving Predictors | Contrast | Estimate (%) | 95% CI | SE | FDR q |
|---|---|---|---|---|---|---|---|---|---|---|
| During-iPBM | Theta | Increase | 6 | Energy, Wavelength, Frequency, Sex, NasalDistance | | | | | | |
| During-iPBM | Theta | Decrease | 79 | Energy, Wavelength, Frequency, Sex, NasalDistance | | | | | | |
| During-iPBM | Beta | Increase | 42 | Energy, Wavelength, Frequency, Sex, NasalDistance | | | | | | |
| During-iPBM | Beta | Decrease | 75 | Energy, Wavelength, Frequency, Sex, NasalDistance | | | | | | |
| During-iPBM | Gamma | Increase | 48 | Energy, Wavelength, Frequency, Sex, NasalDistance | | | | | | |
| During-iPBM | Gamma | Decrease | 74 | Energy, Wavelength, Frequency, Sex, NasalDistance | | | | | | |
| Post-iPBM | Theta | Decrease | 156 | Energy, Wavelength, Frequency, Sex, NasalDistance | | | | | | |
| Post-iPBM | Beta | Increase | 32 | Energy, Wavelength, Frequency, Sex, NasalDistance | | | | | | |
| Post-iPBM | Beta | Decrease | 78 | Energy, Wavelength, Frequency, Sex, NasalDistance | | | | | | |
| Post-iPBM | Gamma | Increase | 43 | Energy, Wavelength, Frequency, Sex, NasalDistance | | | | | | |
| Post-iPBM | Gamma | Decrease | 66 | Energy, Wavelength, Frequency, Sex, NasalDistance | | | | | | |

As shown in **Table S2**, LME results revealed no significant effects of iPBM stimulation parameters or biological factors.

### S5 Model A: Backward elimination trace

**Table S3. Backward elimination trace for the primary linear mixed-effects model (Model A, iPBM).** At each step, Benjamini-Hochberg FDR correction (q=0.05) was applied simultaneously to all fixed-effect p-values in the current model. The predictor whose contrasts yielded the highest FDR-corrected q-value was removed if that value exceeded 0.05. Elimination continued until all remaining predictors were FDR-significant or no predictors remained. Each row documents one removal step for a given frequency band, time window, and cluster ROI combination. The “FDR q-value” column reports the worst (highest) FDR-corrected q-value among the removed predictor's contrasts at that step. Predictor abbreviations: Energy = irradiance level (Low: 5, Mid: 7, High: 9 mW/cm$^2$); Wavelength = 808 or 1064 nm; Frequency = pulsation frequency (10 or 40 Hz); Sex = Male or Female; NasalDistance = z-scored distance.

| Band | Time Window | ROI | Step | Removed Predictor | FDR q | Predictors Before Removal | Predictors After Removal |
|---|---|---|---|---|---|---|---|
| Theta | During-iPBM | Increase | 1 | Wavelength | 0.8468 | Energy, Wavelength, Frequency, Sex, NasalDistance | Energy, Frequency, Sex, NasalDistance |
| Theta | During-iPBM | Increase | 2 | Frequency | 0.6259 | Energy, Frequency, Sex, NasalDistance | Energy, Sex, NasalDistance |
| Theta | During-iPBM | Increase | 3 | Sex | 0.3969 | Energy, Sex, NasalDistance | Energy, NasalDistance |
| Theta | During-iPBM | Increase | 4 | Energy | 0.1834 | Energy, NasalDistance | NasalDistance |
| Theta | During-iPBM | Increase | 5 | NasalDistance | 0.1472 | NasalDistance | |
| Theta | During-iPBM | Decrease | 1 | Sex | 0.9389 | Energy, Wavelength, Frequency, Sex, NasalDistance | Energy, Wavelength, Frequency, NasalDistance |
| Theta | During-iPBM | Decrease | 2 | Energy | 0.6161 | Energy, Wavelength, Frequency, NasalDistance | Wavelength, Frequency, NasalDistance |
| Theta | During-iPBM | Decrease | 3 | Wavelength | 0.6192 | Wavelength, Frequency, NasalDistance | Frequency, NasalDistance |
| Theta | During-iPBM | Decrease | 4 | NasalDistance | 0.6193 | Frequency, NasalDistance | Frequency |
| Theta | During-iPBM | Decrease | 5 | Frequency | 0.0758 | Frequency | |
| Beta | During-iPBM | Increase | 1 | Energy | 0.9108 | Energy, Wavelength, Frequency, Sex, NasalDistance | Wavelength, Frequency, Sex, NasalDistance |
| Beta | During-iPBM | Increase | 2 | Wavelength | 0.9102 | Wavelength, Frequency, Sex, NasalDistance | Frequency, Sex, NasalDistance |
| Beta | During-iPBM | Increase | 3 | Frequency | 0.9102 | Frequency, Sex, NasalDistance | Sex, NasalDistance |

| Band | Time Window | ROI | Step | Removed Predictor | FDR q | Predictors Before Removal | Predictors After Removal |
|---|---|---|---|---|---|---|---|
| Beta | During-iPBM | Increase | 4 | Sex | 0.9152 | Sex, NasalDistance | NasalDistance |
| Beta | During-iPBM | Increase | 5 | NasalDistance | 0.8046 | NasalDistance | |
| Beta | During-iPBM | Decrease | 1 | NasalDistance | 0.572 | Energy, Wavelength, Frequency, Sex, NasalDistance | Energy, Wavelength, Frequency, Sex |
| Beta | During-iPBM | Decrease | 2 | Frequency | 0.2731 | Energy, Wavelength, Frequency, Sex | Energy, Wavelength, Sex |
| Beta | During-iPBM | Decrease | 3 | Sex | 0.2075 | Energy, Wavelength, Sex | Energy, Wavelength |
| Beta | During-iPBM | Decrease | 4 | Energy | 0.1096 | Energy, Wavelength | Wavelength |
| Beta | During-iPBM | Decrease | 5 | Wavelength | 0.1166 | Wavelength | |
| Gamma | During-iPBM | Increase | 1 | Energy | 0.8982 | Energy, Wavelength, Frequency, Sex, NasalDistance | Wavelength, Frequency, Sex, NasalDistance |
| Gamma | During-iPBM | Increase | 2 | Wavelength | 0.8774 | Wavelength, Frequency, Sex, NasalDistance | Frequency, Sex, NasalDistance |
| Gamma | During-iPBM | Increase | 3 | Frequency | 0.8774 | Frequency, Sex, NasalDistance | Sex, NasalDistance |
| Gamma | During-iPBM | Increase | 4 | Sex | 0.8926 | Sex, NasalDistance | NasalDistance |
| Gamma | During-iPBM | Increase | 5 | NasalDistance | 0.9291 | NasalDistance | |
| Gamma | During-iPBM | Decrease | 1 | Energy | 0.8957 | Energy, Wavelength, Frequency, Sex, NasalDistance | Wavelength, Frequency, Sex, NasalDistance |
| Gamma | During-iPBM | Decrease | 2 | NasalDistance | 0.9615 | Wavelength, Frequency, Sex, NasalDistance | Wavelength, Frequency, Sex |
| Gamma | During-iPBM | Decrease | 3 | Frequency | 0.5151 | Wavelength, Frequency, Sex | Wavelength, Sex |
| Gamma | During-iPBM | Decrease | 4 | Wavelength | 0.2783 | Wavelength, Sex | Sex |
| Gamma | During-iPBM | Decrease | 5 | Sex | 0.1574 | Sex | |
| Theta | Post-iPBM | Decrease | 1 | Energy | 0.9645 | Energy, Wavelength, Frequency, Sex, NasalDistance | Wavelength, Frequency, Sex, NasalDistance |
| Theta | Post-iPBM | Decrease | 2 | Wavelength | 0.8228 | Wavelength, Frequency, Sex, NasalDistance | Frequency, Sex, NasalDistance |
| Theta | Post-iPBM | Decrease | 3 | NasalDistance | 0.8228 | Frequency, Sex, NasalDistance | Frequency, Sex |

| Band | Time Window | ROI | Step | Removed Predictor | FDR q | Predictors Before Removal | Predictors After Removal |
|---|---|---|---|---|---|---|---|
| Theta | Post-iPBM | Decrease | 4 | Frequency | 0.4498 | Frequency, Sex | Sex |
| Theta | Post-iPBM | Decrease | 5 | Sex | 0.2391 | Sex | |
| Beta | Post-iPBM | Increase | 1 | Energy | 0.7416 | Energy, Wavelength, Frequency, Sex, NasalDistance | Wavelength, Frequency, Sex, NasalDistance |
| Beta | Post-iPBM | Increase | 2 | Frequency | 0.6982 | Wavelength, Frequency, Sex, NasalDistance | Wavelength, Sex, NasalDistance |
| Beta | Post-iPBM | Increase | 3 | NasalDistance | 0.6901 | Wavelength, Sex, NasalDistance | Wavelength, Sex |
| Beta | Post-iPBM | Increase | 4 | Wavelength | 0.3615 | Wavelength, Sex | Sex |
| Beta | Post-iPBM | Increase | 5 | Sex | 0.3615 | Sex | |
| Beta | Post-iPBM | Decrease | 1 | Frequency | 0.7501 | Energy, Wavelength, Frequency, Sex, NasalDistance | Energy, Wavelength, Sex, NasalDistance |
| Beta | Post-iPBM | Decrease | 2 | Energy | 0.4678 | Energy, Wavelength, Sex, NasalDistance | Wavelength, Sex, NasalDistance |
| Beta | Post-iPBM | Decrease | 3 | Wavelength | 0.5088 | Wavelength, Sex, NasalDistance | Sex, NasalDistance |
| Beta | Post-iPBM | Decrease | 4 | Sex | 0.5088 | Sex, NasalDistance | NasalDistance |
| Beta | Post-iPBM | Decrease | 5 | NasalDistance | 0.7202 | NasalDistance | |
| Gamma | Post-iPBM | Increase | 1 | Energy | 0.8965 | Energy, Wavelength, Frequency, Sex, NasalDistance | Wavelength, Frequency, Sex, NasalDistance |
| Gamma | Post-iPBM | Increase | 2 | NasalDistance | 0.8817 | Wavelength, Frequency, Sex, NasalDistance | Wavelength, Frequency, Sex |
| Gamma | Post-iPBM | Increase | 3 | Wavelength | 0.3342 | Wavelength, Frequency, Sex | Frequency, Sex |
| Gamma | Post-iPBM | Increase | 4 | Frequency | 0.3342 | Frequency, Sex | Sex |
| Gamma | Post-iPBM | Increase | 5 | Sex | 0.3125 | Sex | |
| Gamma | Post-iPBM | Decrease | 1 | NasalDistance | 0.9881 | Energy, Wavelength, Frequency, Sex, NasalDistance | Energy, Wavelength, Frequency, Sex |
| Gamma | Post-iPBM | Decrease | 2 | Energy | 0.6974 | Energy, Wavelength, Frequency, Sex | Wavelength, Frequency, Sex |
| Gamma | Post-iPBM | Decrease | 3 | Frequency | 0.6936 | Wavelength, Frequency, Sex | Wavelength, Sex |

| Band | Time Window | ROI | Step | Removed Predictor | FDR q | Predictors Before Removal | Predictors After Removal |
|---|---|---|---|---|---|---|---|
| Gamma | Post-iPBM | Decrease | 4 | Wavelength | 0.2761 | Wavelength, Sex | Sex |
| Gamma | Post-iPBM | Decrease | 5 | Sex | 0.1415 | Sex | |

### S6 Model B: Carry-over analysis

**Table S4. Results of the carry-over analysis (Model B).** This model tested whether the energy level of the immediately preceding recording session (PrevEnergyLevel; reference: Low) predicted the current session's EEG band power response, with subject as a random intercept. Backward elimination with FDR correction was applied identically to Model A. Because tPBM and iPBM recordings were alternated within each session, preceding sessions include both tPBM and iPBM doses. Results are shown for both tPBM and iPBM outcomes to provide a complete picture of potential cross-modality carry-over effects.

| Modality | Time Window | Band | ROI | N Electrodes | Contrast | Estimate (%) | 95% CI | SE | FDR q |
|---|---|---|---|---|---|---|---|---|---|
| tPBM | During-tPBM | Theta | Decrease | 119 | | | | | |
| tPBM | During-tPBM | Alpha | Decrease | 172 | | | | | |
| tPBM | During-tPBM | Beta | Increase | 23 | | | | | |
| tPBM | During-tPBM | Beta | Decrease | 124 | | | | | |
| tPBM | During-tPBM | Gamma | Increase | 42 | | | | | |
| tPBM | During-tPBM | Gamma | Decrease | 68 | | | | | |
| tPBM | Post-tPBM | Theta | Decrease | 180 | | | | | |
| tPBM | Post-tPBM | Alpha | Decrease | 161 | | | | | |
| tPBM | Post-tPBM | Beta | Decrease | 120 | | | | | |
| tPBM | Post-tPBM | Gamma | Increase | 45 | | | | | |

| Modality | Time Window | Band | ROI | N Electrodes | Contrast | Estimate (%) | 95% CI | SE | FDR q |
|---|---|---|---|---|---|---|---|---|---|
| iPBM | During-iPBM | Theta | Increase | 6 | | | | | |
| iPBM | During-iPBM | Theta | Decrease | 79 | | | | | |
| iPBM | During-iPBM | Beta | Increase | 42 | | | | | |
| iPBM | During-iPBM | Beta | Decrease | 75 | | | | | |
| iPBM | During-iPBM | Gamma | Increase | 48 | | | | | |
| iPBM | During-iPBM | Gamma | Decrease | 74 | Previous High vs Low dose | -7.804 | [-14.626, -0.982] | 3.481 | 0.0250* |
| iPBM | During-iPBM | Gamma | Decrease | 74 | Previous Mid vs Low dose | -9.116 | [-15.903, -2.328] | 3.463 | 0.0170* |
| iPBM | Post-iPBM | Theta | Decrease | 156 | | | | | |
| iPBM | Post-iPBM | Beta | Increase | 32 | | | | | |
| iPBM | Post-iPBM | Beta | Decrease | 78 | | | | | |
| iPBM | Post-iPBM | Gamma | Increase | 43 | | | | | |
| iPBM | Post-iPBM | Gamma | Decrease | 66 | | | | | |

As shown in **Table S4**, For tPBM outcomes, no significant carry-over effects were detected in any band, phase, or ROI direction. For iPBM outcomes, significant carry-over effects were detected only in the gamma band during stimulation within the negative (power decrease) ROI: preceding sessions with Mid or High energy levels were associated with significantly more pronounced gamma suppression compared to Low energy.

### S7 Energy-efficiency analysis

**Table S5. Paired comparison of the energy efficiency of iPBM versus tPBM in modulating EEG band power.** Efficiency was defined as the mean percent change in band power from the PRE period, averaged across the electrodes of the cluster ROI, divided by the delivered energy of that recording (fluence, $J/cm^2$); units are therefore percent change per $J/cm^2$. Efficiency was computed separately for each recording and then averaged across the four recordings per modality to yield one value per subject per modality. Each modality was evaluated within its own FWER-significant spatiotemporal cluster ROI (**Section 2.4.3.1**), separated into Increase and Decrease sub-ROIs according to the sign of the group-mean percent change across cluster electrodes. A band × time window × ROI combination was compared only when both modalities contributed at least three cluster electrodes, which excluded the alpha band and yielded the eight comparisons listed. Median tPBM Eff and Median iPBM Eff are the across-subject medians of these signed subject-level efficiencies, so negative values denote power decreases per unit delivered energy. The Median |iPBM/tPBM| Ratio is the median of the within-subject ratio of absolute efficiencies rather than the ratio of the two medians; values above one indicate a larger neural effect per joule for iPBM. Each combination was tested with a two-sided Wilcoxon signed-rank test on the paired subject-level efficiencies (n=46), and FDR correction (q=0.05) was applied across all eight tests. Significant indicates q<0.05.

| Band | Time Window | ROI | N Subjects | Median tPBM Eff | Median iPBM Eff | Median \|iPBM/tPBM\| Ratio | Wilcoxon Statistic | p | FDR q | Significant |
|---|---|---|---|---|---|---|---|---|---|---|
| Theta | DURING | Decrease | 46 | -0.1341 | -1.0674 | 14.74 | 419 | 0.188012 | 0.188012 | FALSE |
| Theta | POST | Decrease | 46 | -0.3051 | -6.7807 | 18 | 318 | 0.014294 | 0.016336 | TRUE |
| Beta | DURING | Increase | 46 | 0.2624 | 4.8856 | 16.44 | 123 | 1.00E-06 | 4.00E-06 | TRUE |
| Beta | DURING | Decrease | 46 | -0.1841 | -3.7259 | 22.42 | 190 | 6.30E-05 | 0.000126 | TRUE |
| Beta | POST | Decrease | 46 | -0.3015 | -7.7954 | 17.14 | 272 | 0.002817 | 0.003756 | TRUE |
| Gamma | DURING | Increase | 46 | 0.6883 | 13.5217 | 14.34 | 109 | 0 | 0 | TRUE |
| Gamma | DURING | Decrease | 46 | -0.1531 | -4.2329 | 32.4 | 259 | 0.001676 | 0.002682 | TRUE |
| Gamma | POST | Increase | 46 | 1.7388 | 32.4527 | 14.53 | 181 | 3.80E-05 | 0.000101 | TRUE |
